# Genomic plasticity is a blueprint of diversity in *Salmonella* lineages

**DOI:** 10.1101/2023.12.02.569618

**Authors:** Simran Krishnakant Kushwaha, Abhirath Anand, Yi Wu, Hugo Leonardo Ávila, Thomas Sicheritz-Pontén, Andrew Millard, Sandhya Amol Marathe, Franklin L. Nobrega

## Abstract

The flexible genome, organised in regions of genomic plasticity (RGP), serves as a potent facilitator of the dynamic evolution of bacterial genomes through gene acquisition and loss. Here we explored the genomic plasticity across *Salmonella* lineages, revealing a purposeful, non-random integration pattern of pathogenicity-related gene clusters into specific RGP. Noteworthy examples include the correlation between the type I-E CRISPR-Cas system, gold tolerance, and specific RGP. The scattered prevalence of RGP across *Salmonella* lineages profoundly shapes the pathogenicity makeup of *Salmonella* strains. The preferences of RGP seem guided by conserved flanking genes that likely share regulatory and functional coordination. For example, metal resistance genes are predominant in RGP positioned near stress resistance genes, indicating a regulatory network to efficiently counter stressors. Additionally, we observed that different plasmid incompatibility types and prophage genera carry distinct pathogenicity genes. Similar to RGP, their distribution across *Salmonella* lineages plays a critical role in defining pathogenicity. By uncovering these intricate connections among gene clusters, RGP, mobile genetic elements, and pathogenic attributes, our study offers novel insights into the evolutionary trajectory of *Salmonella* and holds promise for predicting future adaptations and developing targeted interventions against infections.

## Introduction

The interplay between conserved and variable features in bacterial genomes plays a crucial role in shaping the diversity and adaptability of different species^1^. Within a species, the core genome, comprising genes universally present, handles essential cellular functions. In contrast, the flexible genome consists of genes that vary between individual strains, allowing bacteria to adapt to specific environments and acquire pathogenic traits^2–4^. These variable genes are often organised into regions of genomic plasticity (RGP)^5^, typically associated with mobile genetic elements (MGEs). These elements serve as potent facilitators for acquiring genes related to virulence, antibiotic and stress resistance, and anti-phage immunity, contributing to the dynamic evolution of the bacterial genome^6–9^. Exploring this genomic plasticity is crucial for understanding bacterial evolution, phylogeny, and pathogenic potential.

*Salmonella* offers an excellent model for studying these variable genomic features. Its diverse spectrum of species, subspecies, and serovars showcases the inherent flexibility in its genome, a pivotal factor in shaping both the phylogeny and pathogenic potential of *Salmonella*^10–12^. Consequently, exploring the genomic plasticity of *Salmonella* becomes a key avenue for gaining insights into its evolution as a pathogen.

To gain further understanding of the structural and functional features of RGP in *Salmonella*, we carried out a comprehensive analysis of 12,244 *Salmonella* spp. genomes. Our findings revealed that gene clusters associated with virulence, stress resistance, antibiotic resistance, and anti-phage defence exhibit specific preferences for RGP integrated into distinct genomic spots. These preferences seem to be influenced by neighbouring genes that likely share regulatory and functional coordination. The irregular distribution of these genomic spots across diverse *Salmonella* lineages establishes a blueprint for pathogenicity and survival strategies. Deciphering the complex interplay between pathogenicity-related gene clusters and RGP not only improves our understanding of *Salmonella* evolution, but also enables us to uncover novel pathogenicity genes, anticipate future adaptations, and identify targets for disease prevention, management, and therapeutic interventions.

## Results

### The mobilome of *Salmonella* is highly variable across lineages

MGEs play a pivotal role in driving genetic diversity and shaping the evolutionary trajectories of bacteria, enabling them to adapt to various environmental challenges^13^. One significant way through which MGEs exert their influence is by facilitating the horizontal transfer of genes associated with pathogenicity traits, directly impacting the potential of bacterial pathogens. To determine the broad relevance of specific MGEs in defining specific pathogenicity attributes of *Salmonella*, we explored the variation in plasmids and prophages across 12,244 *Salmonella* genomes (**Supplementary Table 1**). Our dataset included representative strains from the two species and six subspecies of *Salmonella*, as well as 46 serovars of *Salmonella enterica* subsp. *enterica* (**Fig. 1a**). These serovars were grouped into host-specific, host-adapted and broad-host range, with those lacking sufficient information categorised within the host range of unknown origin. As expected, the most prevalent serovars were Typhi (2,440 strains) and Typhimurium (2,170 strains), the main causative agents of typhoid fever^14^ and gastroenteritis^15^ in humans, respectively. The genome sequence of these strains was used to infer a phylogenetic topology representing the genomic diversity within the genus *Salmonella* (**Fig. 1b**). The overall topology of the phylogeny is in accordance with the phenogram created previously from concatenated MLST genes of a smaller number of genomes^16^.

**Fig. 1.**
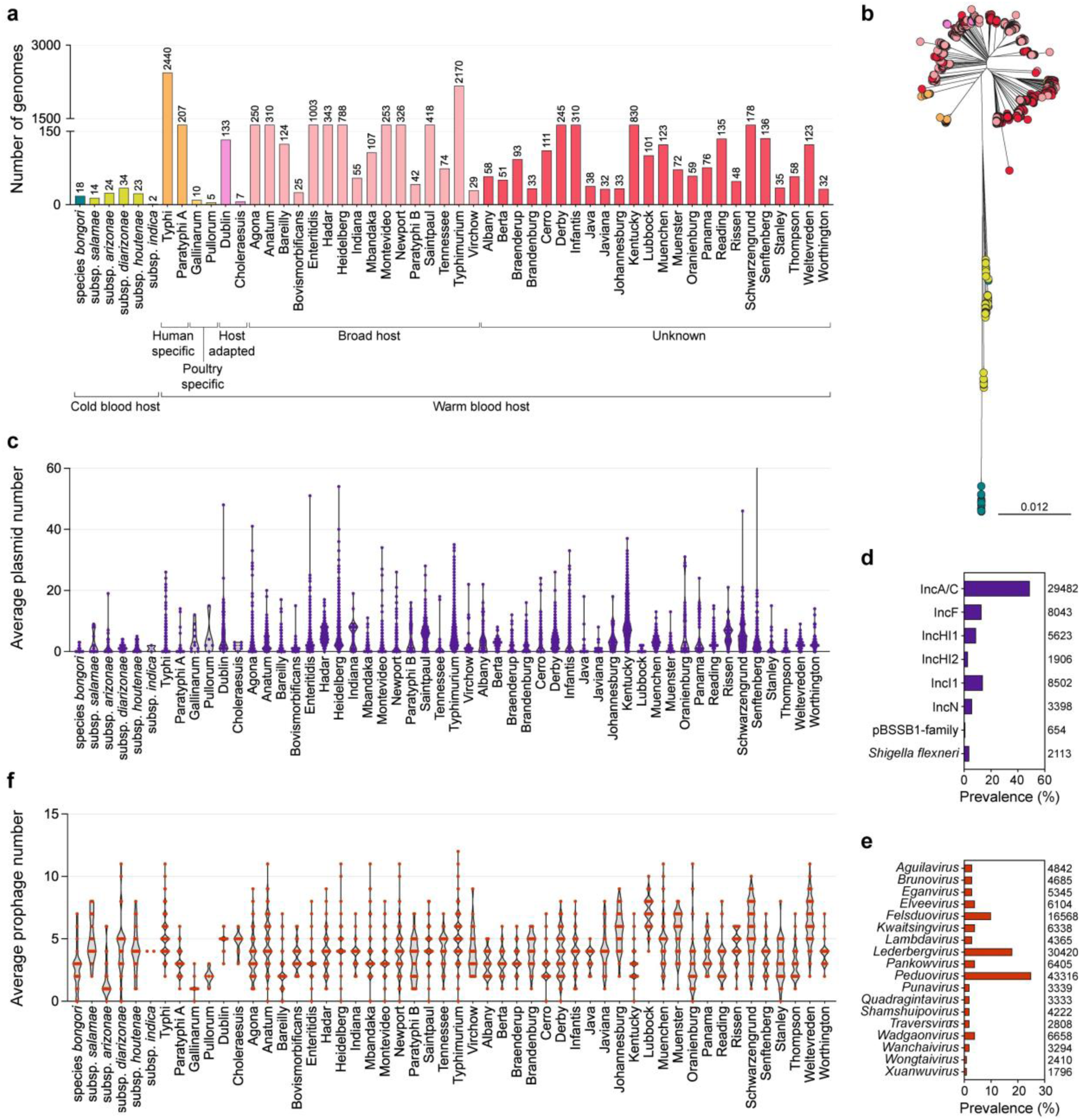
| Characterisation of the *Salmonella* database. **a** Distribution of strains from various *Salmonella* subspecies and serovars, categorised by their host specificity. **b** Phylogenetic distribution of the *Salmonella* strains with a colour scheme analogous to (a). **c** Average plasmid content in *Salmonella* subspecies and serovars. **d** Prevalence of plasmid incompatibility groups in *Salmonella*. The *Shigella flexneri* plasmid incompatibility group refers to virulence plasmid pINV. **e** Average prophage content in *Salmonella* subspecies and serovars. **f** Prevalence of prophage genera in *Salmonella*.

Analysis of plasmid prevalence across different *Salmonella* subspecies and serovars revealed a variable abundance of these plasmid contigs (**Fig. 1c, Supplementary Table 2**). For instance, serovar Kentucky exhibited an average of 10 plasmid contigs, whereas most strains of serovar Paratyphi A lacked any plasmid contigs. Among the identified plasmids, the most prevalent were those belonging to the IncA/C group (39%, 29,482), IncI group (11%, 8,502), and IncF group (11%, 8,043) (**Fig. 1d**, **Supplementary Table 3**). Notably, IncA/C plasmids were predominantly found (58%, 17,091) in serovar Typhimurium, while IncHI1 (40%, 2,244) and IncN (30%, 1,025) were more commonly observed in serovar Typhi (**Supplementary Table 3**). Other plasmid types exhibited a more even distribution across different species.

Analysis of prophage prevalence in *Salmonella* shows that the vast majority of strains (96.6%, 11,829) carry at least one prophage, accounting for a total of 52,555 prophage regions (**Supplementary Table 4**). From this total, the taxonomy of 18,785 complete dsDNA prophage regions was determined using taxmyPHAGE (https://github.com/amillard/tax_myPHAGE). In most cases, we identified prophage regions associated with phages from multiple families, genera and species, resulting in a total of 172,862 entries. All these phages belong to the kingdom *Heunggongvirae*, phylum *Uroviricota*, and class *Caudoviricetes*. Within *Caudoviricetes*, 25% (43,316) of the phage regions belong to the genus *Peduovirus*, 18% (30,402) to the genus *Lederbergvirus* and 10% (16,568) to the genus *Felsduovirus* (**Fig. 1e**). The remaining phages are distributed across 68 other identified genera, though in smaller quantities. The most commonly identified phages include *Lederbergvirus Salmonella* phage BTP1, SE1Spa, P22, ST64T, *Enterobacteria* phage HK620 and *Shigella* phage Sf6. Similar to plasmids, the average number of prophages per strain varies among serovars, with serovar Lubbock averaging seven prophages, whereas serovar Gallinarum has only one (**Fig. 1f**).

In summary, our findings highlight the remarkable diversity observed in the mobilome of *Salmonella*. This diversity is reflected in the abundance and types of MGEs present per species, subspecies, and serovars. The variable nature of the mobilome and the resulting diversity in gene composition are expected to play a critical role in shaping the pathogenicity, adaptation, and distribution of *Salmonella*.

### Virulence determinants are more prevalent in chromosomal regions

We next analysed the presence of factors contributing to the survival and adaptation of *Salmonella* to the environment. These included a set of virulence factors, antibiotic resistance genes, stress response genes, and phage-resistance genes (i.e., anti-phage defence systems) (the complete list of genes can be found in **Supplementary Table 5**). This analysis revealed the presence of virulence factors predominantly in *S. enterica* subsp. *enterica*, with an average of 46 virulence factors per strain (**Fig. 2a**, **Supplementary Tables 6 and 7**). Interestingly, *S. bongori* has the lowest number of virulence genes, 20. In comparison to most *S. enterica* subsp. *enterica* strains, *S. bongori* lacks the *Salmonella* pathogenicity island 2 (SPI-2), which encodes a type III secretion system (T3SS) that plays a central role in systemic infections and the intracellular phenotype of *S. enterica*, except for one strain (accession no. 1173775.3) that also groups with cold-blooded subspecies of *S. enterica* in the phylogenetic tree (**Fig. 1b**). The ability of SPI-2 to transfer to, and be functional in, *S. bongori* has been previously demonstrated experimentally^17^ but, to our knowledge, not yet found in nature^16^. *S. bongori* do contain the SPI-1 with a T3SS that promotes invasion of epithelial cells through the secretion of different effector proteins^18^. Curiously, SPI-1 is prevalent across all *Salmonella* species, subspecies and serovars, but with variations in the presence of secreted effectors encoded by *spt* and *slr*, as well as *avr* and *ssp* genes, especially the latter (**Supplementary** Fig. 1).

**Fig. 2.**
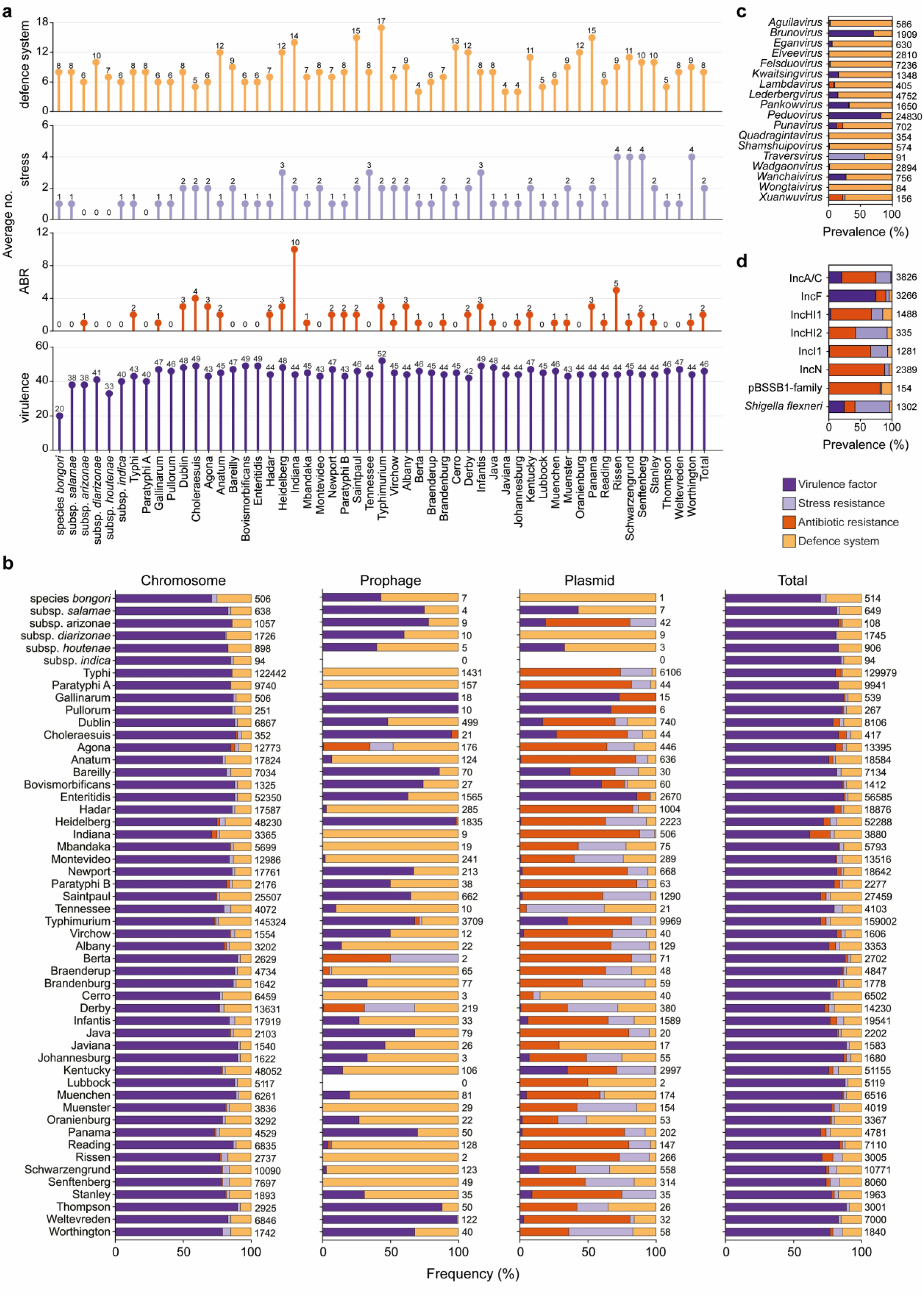
| Prevalence and distribution of pathogenicity determinants in *Salmonella*. **a** Prevalence of virulence factors, stress resistance genes, antibiotic resistance genes, and anti-phage defence systems across *Salmonella* subspecies and serovars. **b** Distribution of the pathogenicity determinants across chromosome, prophages, and plasmids. **c** Distribution of the pathogenicity determinants across prophage genera present in *Salmonella* at >1% abundance. **d** Distribution of the pathogenicity determinants across plasmid incompatibility groups present in *Salmonella*. In all panels, virulence factors, stress resistance genes, antibiotic resistance genes, and defence systems are coloured according to the key.

Poultry-host-specific serovars Gallinarum and Pullorum share a more recent common ancestor with the broad host range serovar Enteritidis, but the gene *rck*, contributed by *Salmonella* virulence plasmid and responsible for evading the host immune response and surviving inside the host^19^, is absent in Gallinarum and Pullorum (**Supplementary** Fig. 1). This gene is likely involved in the broader host range of Enteritidis strains.

Our analysis revealed that the majority of virulence factors are in chromosomal regions (**Fig. 2b, Supplementary** Fig. 1, **Supplementary Table 6**). However, certain virulence genes are more commonly associated with prophages or plasmids. For example, genes *sod* and *grv,* critical to the bacterial response to oxidative stress and their ability to survive within immune cells^20,21^, are frequently found in prophage regions, mostly of *Peduovirus* (80.1% *sod*, 86.1% *grv*) and *Brunovirus* (3.7% *sod*, 3.7% *grv*) genera (**Supplementary** Fig. 1). These prophages seem to preferentially carry virulence factors (**Fig. 2c**). Surprisingly, in contrast to existing literature^22^, we found that the majority of *gog* genes, which are associated with an anti-inflammatory function^23^, are located on the chromosome (1,574 out of 1,703) rather than a prophage region (**Supplementary** Fig. 1**, Supplementary Table 6**). The well-known virulence genes *spv* (involved in intracellular survival and evasion of the host immune response^24^), *pef* (plasmid encoded fimbriae, important for colonisation of the host and establishment of infection^25,26^), and *rck* (contributing to evasion of the host immune response^27^) are predominantly found on plasmids (**Supplementary** Fig. 1), particularly those belonging to the IncF group (**Fig. 2d, Supplementary Table 8**). These *Salmonella* virulence plasmids (pSV) containing the *spv* genes were identified in *S. enterica* subsp. *arizonae* and *S. enterica* subp. *enterica* serovar Typhimurium, Dublin, Enteritidis, Choleraesuis, Gallinarum, and Pullorum, consistent with the existing literature^28–31^. Genes *pef* and *rck* have also been reported previously in pSV^32^. The *fyu* and *ybt* genes involved in iron acquisition^33^ were predominantly associated with IncA/C plasmids. Notably, we did not find any virulence factors on IncHI2, IncN, and pBSSB1-family plasmids (**Fig. 2d**, **Supplementary Table 8**).

In summary, our analysis reveals that virulence factors in *Salmonella* are primarily found within chromosomal regions, but specific gene clusters are preferentially located in prophages of the *Brunovirus* and *Peduovirus* genera, and plasmids of the IncF group.

### Antibiotic resistance genes are primarily located within plasmids

Plasmids serve as the primary reservoir for antibiotic resistance (ABR) genes in *Salmonella* (**Fig. 2b**). Specifically, 78% of the ABR genes identified in *Salmonella* were located on plasmids, with 21% found on chromosomal regions (predominantly streptothricin and fosfomycin), and 1% associated with prophages (mostly of the *Xuanvirus* genus, **Fig. 2c**). While ABR levels in *Salmonella* prophages are lower compared to those in plasmids or the chromosome, they surpass previously reported general prophage analyses^34,35^. Most ABR genes were found across multiple Inc plasmids schemes, but majorly in IncN and IncA/C (**Fig. 2d, Supplementary Table 8**).

Consistently, the strains exhibited the highest resistance to tetracycline, streptomycin, sulphonamide, and beta-lactam antibiotics (**Supplementary** Fig. 1), which aligns with previous reports^36^. Importantly, ABR genes were prevalent in strains of *S. enterica* subsp. *enterica* but negligible (≤ 1 ABR gene) in other *S. enterica* subspecies (except *S. enterica* subsp. *arizonae*) and *S. bongori*. This pattern does not seem to be strongly driven by a lower abundance of plasmids (r^2^ = 0.2965, p = <0.0001, **Supplementary** Fig. 2a). Notably, serovar Paratyphi A showed minimal presence of ABR genes, as plasmids are also almost absent in this serovar. On the other hand, serovars Indiana and Rissen carried an average of 10 and 5 ABR genes, respectively, indicating multidrug resistance, consistent with previous reports^37,38^ (**Supplementary** Fig. 1, **Supplementary Tables 5, 6 and 7)**. The serovars Heidelberg, Typhimurium, Newport, and Enteritidis are known to cause the majority of outbreaks, and 89%, 75%, 32% and 10% of strains from these serovars contain ABR genes (**Supplementary** Fig. 1**, Supplementary Table 6**). Importantly, the presence of resistance to colistin, an antibiotic of last resort, was detected in 2.4% (288) of strains belonging to *S. enterica* subsp. *enterica*, with a predominant occurrence in serovars Saintpaul, Cholerasuis, and Paratyphi B (**Supplementary** Fig. 1).

In summary, our results reinforce the role of plasmids in influencing antibiotic resistance patterns in *Salmonella*, and highlight plasmids of all schemes are drivers of ABR dissemination. Moreover, the higher prevalence of antibiotic resistance in *S. enterica* subsp. *enterica*, as compared to other *Salmonella* species and subspecies, underscores the influence of human antibiotic usage in promoting the spread of antibiotic resistance.

### Stress-resistance genes are primarily located on plasmids and chromosomal regions

The presence of acid, biocide, and heavy metal resistance genes is closely linked to the maintenance and spread of antimicrobial resistance^39–41^. Interestingly, we observed that the two multidrug resistant serovars, Indiana and Rissen, exhibit the highest prevalence of *qac* genes, which are small multidrug resistance efflux proteins associated with increased tolerance to quaternary ammonium compounds (QAC) and other cationic biocides^42^ (**Supplementary** Fig. 1). *qac* genes are generally found in MGEs; here, they were found on IncI1 and IncA/C plasmids in over 75% of the cases (**Supplementary Table 8**).

Curiously, most strains in our dataset do not carry any heat-resistant genes, except for a small percentage (<20%) of strains from serovars Montevideo, Senftenberg, and Worthington, and the majority of these genes are located on IncA/C plasmids. On the other hand, *Salmonella* strains commonly exhibited resistance to heavy metals, with approximately 80% of the strains carrying genes conferring resistance to gold (**Supplementary** Fig. 1). The only exceptions are *S. enterica* subsp. *houtenae* and *S. enterica* subsp. *enterica* serovars Typhi and Paratyphi A, which do not carry the *gol* cluster responsible for gold resistance. Serovars Heidelberg and Infantis show a high incidence (>95%) of arsenic resistance genes (*ars*), while serovars Tennessee, Rissen, Schwarzengrund, Worthington, and Senftenberg exhibit frequent (>80%) copper (*pco*) and silver (*sil*) resistance (**Supplementary** Fig. 1, **Supplementary Table 6**).

Curiously, genes that confer resistance to heavy metals such as gold, arsenic, copper, and silver are predominantly located in chromosomal regions, while those associated with mercury and tellurite resistance are commonly found on plasmids (IncA/C, and IncHI1 and IncHI2, respectively) (**Supplementary** Fig. 1). Among the different plasmid schemes, *Shigella flexneri* (virulence plasmid pINV^43^) and IncHI2 plasmids are those most frequently associated with stress resistance genes, but IncA/C plasmids, due to their abundance, are responsible for the movement of most stress resistance genes (**Fig. 2d**, **Supplementary Table 8**).

In summary, our findings reveal that genes associated with resistance to biocides, heat, and heavy metals such as mercury and tellurite are primarily found on plasmids, while resistance to gold, arsenic, copper, and silver is commonly found within chromosomal regions. The elevated levels of heavy metal resistance observed in specific serovars raise concerns about the use of heavy metal-based compounds in animal-production settings.

### Anti-phage defence systems are more prevalent in chromosomal regions

Anti-phage defence systems were found to be prevalent among *Salmonella* strains, with an average of eight defence systems per strain. This is higher than the average found in *Escherichia coli* (six)^44^ or *Pseudomonas aeruginosa* (seven)^45^ in previous studies. However, there is considerable variation in the number of defence systems carried by different subspecies and serovars. For example, serovars Typhimurium (17), Saintpaul (15), Panama (15), and Indiana (14) exhibit the highest prevalence of defence systems, while serovars Berta, Javiana, and Johannesburg have the lowest (4) (**Fig. 2a**, **Supplementary Tables 6 and 7**).

Among the 90 defence system subtypes identified in *Salmonella* strains, the most prevalent were the restriction-modification (RM) and type I-E CRISPR-Cas systems, which are present in almost all subspecies and serovars (**Supplementary** Fig. 1). However, the CRISPR-Cas system is absent in serovars Brandenburg, Lubbock, and Worthington. We noted significant variation in the prevalence of other defence systems across the *Salmonella* genus (**Supplementary** Fig. 1). Notably, each serovar appears to have a distinct profile of defence systems, suggesting selection of the most beneficial systems in specific environments or host interactions, as previously observed for distinct *E. coli* phylogroups^44^. For example, in serovars Typhi and Paratyphi A, the 3HP and Druantia type III systems are highly abundant. On the other hand, in Typhimurium, we observed a predominance of the BREX type I, Mokosh type II, PARIS types I and II, and Retron II-A defence systems. Strains of Enteritidis exhibit an enrichment in CBASS type I, while Gallinarium and Pullorum frequently harbour Mokosh type I in addition to CBASS type I. Additionally, we found specific defence systems enriched in particular species and subspecies. For instance, dCTP deaminase is more prevalent in *S. bongori*, Septu type I in *S. enterica* subsp. *indica* and *salamae*, and Gabija in *S. enterica* subsp. *arizonae* (**Supplementary** Fig. 1).

In general, defence systems, including the abundant RM and CRISPR-Cas systems, are more frequently found within chromosomal regions (94%, **Fig. 2b**) compared to prophages or plasmids. However, prophages of all genera except *Brunovirus*, *Peduovirus* and *Traversvirus* show a clear preference for carrying defence systems over other types of pathogenicity-related genes (**Fig. 2c**), and the defence systems 3HP, AbiL, BstA, Kiwa, Retron types I-A, I-C, and VI are predominantly found on prophages (**Supplementary** Fig. 1 and 2b). Other defence systems, such as AbiQ, Bunzi, Gao_19, Lit, PifA, ppl, retron type V, SoFic, and tmn are frequently associated with plasmids. Among these, the systems ppl (89%) and GAO_19 (64%) are primarily linked to IncA/C plasmids, while pifA is mostly found (58%) on IncI1 plasmids. Lit (100%) and Bunzi (49%) are often identified on IncHI plasmids (**Supplementary** Fig. 2c, **Supplementary Table 8**). Interestingly, although plasmids of all types often accommodate a greater abundance of other pathogenicity-related elements (**Fig. 2d**), it is noteworthy that the IncHI1 and pBSSB1-family plasmids demonstrate a higher inclination toward carrying defence systems compared to other plasmid schemes.

In summary, anti-phage defence systems are widespread in *Salmonella*, with a notable prevalence of the R-M and the CRISPR-Cas systems. The significant variation in defence system repertoire across *Salmonella* species and serovars observed here highlights the significance of these systems in the evolution and adaptation of this pathogenic bacterium. This is likely influenced by their differential prevalence in distinct MGEs.

### Gene clusters integrate into preferential spots in the *Salmonella* genome

Our analysis uncovered substantial variability in the presence and arrangement of genes associated with virulence, antibiotic resistance, stress response, and anti-phage defence genes among different *Salmonella* strains. This variability strongly suggests the occurrence of genomic rearrangements involving the insertion and deletion of genes. To gain a deeper understanding of genome plasticity within *Salmonella*, we performed a comprehensive mapping analysis using PPanGGoLiN^46^ and panRGP^47^ to identify RGP.

Our findings show that only 4.6% of the genes are conserved in nearly all *Salmonella* genomes (3,575 persistent genes, among which 65 are core genes), while 5.5% (4,256) were present at intermediate frequencies (shell genes), and ∼90% (69,678) at low frequency (cloud genes) (**Fig. 3a**). Analysis of the average gene length showed that persistent and core genes (873 bp) are significantly longer than shell genes (558 bp) and cloud genes (420 bp) (**Fig. 3b, Supplementary Table 9**). In eukaryotes, longer genes are suggested to be more evolutionarily conserved and associated with important biological processes^48–51^. This observation aligns with our findings in *Salmonella*, as functional analysis of the persistent genes revealed their essential roles in survival and fitness (**Supplementary Table 9**). In contrast, shorter gene length is associated with high expression^52^, providing an advantage in response to stimuli^53^. This observation is consistent with the role of accessory shell and cloud genes, which are likely to confer fitness benefits under specific environmental and stress conditions.

**Fig. 3.**
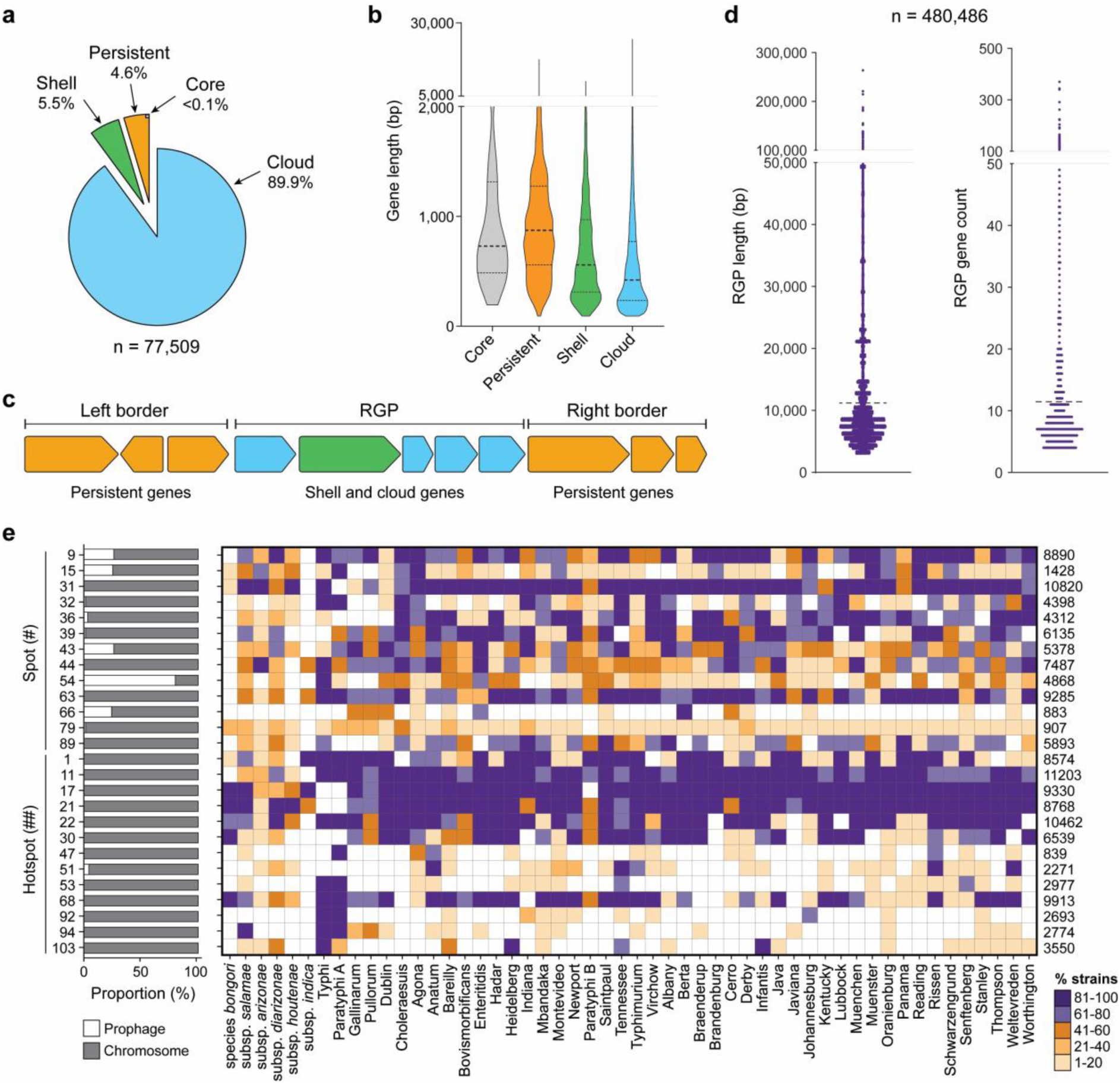
| Pangenome analysis of 12,244 *Salmonella* genomes. **a** Partition of *Salmonella* gene families by PPanGGOLiN, based on their conservation across strains. Core, conserved in all genomes; persistent, conserved in almost all genomes; shell, moderately conserved; cloud, poorly conserved. **b** Length of core, persistent, shell, and cloud genes. **c** Schematic representation of a region of genomic plasticity (RGP), consisting of shell and cloud genes, identified in between conserved border genes. **d** Length and gene count in the identified RGP. **e** Distribution of 26 integration spots across chromosome and prophages, and their prevalence across *Salmonella* subspecies and serovars. The 26 spots correspond to those identified in >1% strains of *Salmonella* containing pathogenicity determinants. Spots (#) are characterised by the presence of > 1 and ≤ 100 unique genes, and hotspots (##) by the presence of > 100 genes.

Clusters of accessory shell and cloud genes form RGP, primarily originating from horizontal gene transfer events. These RGP can be grouped into specific insertion spots based on the presence of conserved flanking persistent genes (**Fig. 3c**). Our analysis identified a total of 673,113 RGP, among which 71.4% (480,486) were clustered in 1,345 spots. These RGP have an average length of 11,182 bp and an average of 11 genes per RGP (**Fig. 3d**). The majority of the RGP (96.5%) is located in the bacterial chromosome and the remaining 3.5% are prophages (i.e. border genes of the spot correspond to those of the prophage) (**Supplementary Table 10**). Out of the 1,345 spots, 74.65% (1,004) were specific to a single type of RGP, while the remaining spots exhibited the potential to harbour a diverse array of RGP families with diverse gene content. Importantly, 1.64% (22) of these spots could harbour >100 distinct RGP families (**Supplementary Table 11)**, suggesting higher rates of gene acquisition and underscoring these regions as hotspots for gene integration^47^.

We screened all spots for the presence of virulence genes, antibiotic resistance genes, stress resistance genes, and defence systems (**Supplementary Table 12**). Among the resulting 266 spots, we selected those with variable content present in at least 1% of the strains, yielding 26 spots (#) (13 of which are hotspots, ##) for further analysis. Some spots were relatively specific to certain serovars, such as spot ##47 in serovars Paratyphi A, Anatum, Agona and Rissen, spot ##92 in Typhi, Paratyphi A, Johannesburg, or spot ##94 in host-specific serovars. In contrast, other spots were widely distributed, such as spots #9 or ##22 (**Fig. 3e**).

When examining the gene content of these spots related to virulence, stress, antibiotic resistance, and defence systems, we can observe that genes with specific functions show a clear propensity to congregate within particular locations (**Fig. 4a**). For instance, *lfp* genes preferentially localise in hotspot ##30, while *fae* genes predominantly localise in spot #36 (**Fig. 4b, Supplementary Table 12**). The gene cluster conferring tolerance to gold (*gol*) distinctly favours spot ##17; the absence of this spot in serovars Typhi and Paratyphi A leads to the absence of gold resistance (**Supplementary** Fig. 1). However, spot ##17 is present in a few strains of *S. enterica* subsp. *houtenae*, where gold resistance is lacking, indicating that the presence of this spot does not consistently correlate with gold resistance. Similarly, the gene cluster conferring arsenic resistance (*ars*) predominantly localises in spot ##103, which is prevalent in serovars Heidelberg and Infantis, the strains of which display the highest prevalence of arsenic resistance. However, spot ##103 is also frequently found in serovar Typhi, where arsenic resistance is absent. Collectively, these findings underscore that the presence of a spot where a specific gene cluster predominantly localises does not unequivocally signify the presence of said gene cluster; conversely, the absence of the site often corresponds to the absence of the specific gene cluster. In cases of the former, other influencing factors may contribute to the selection for the presence of such gene clusters, potentially spurred by environmental pressures.

**Fig. 4.**
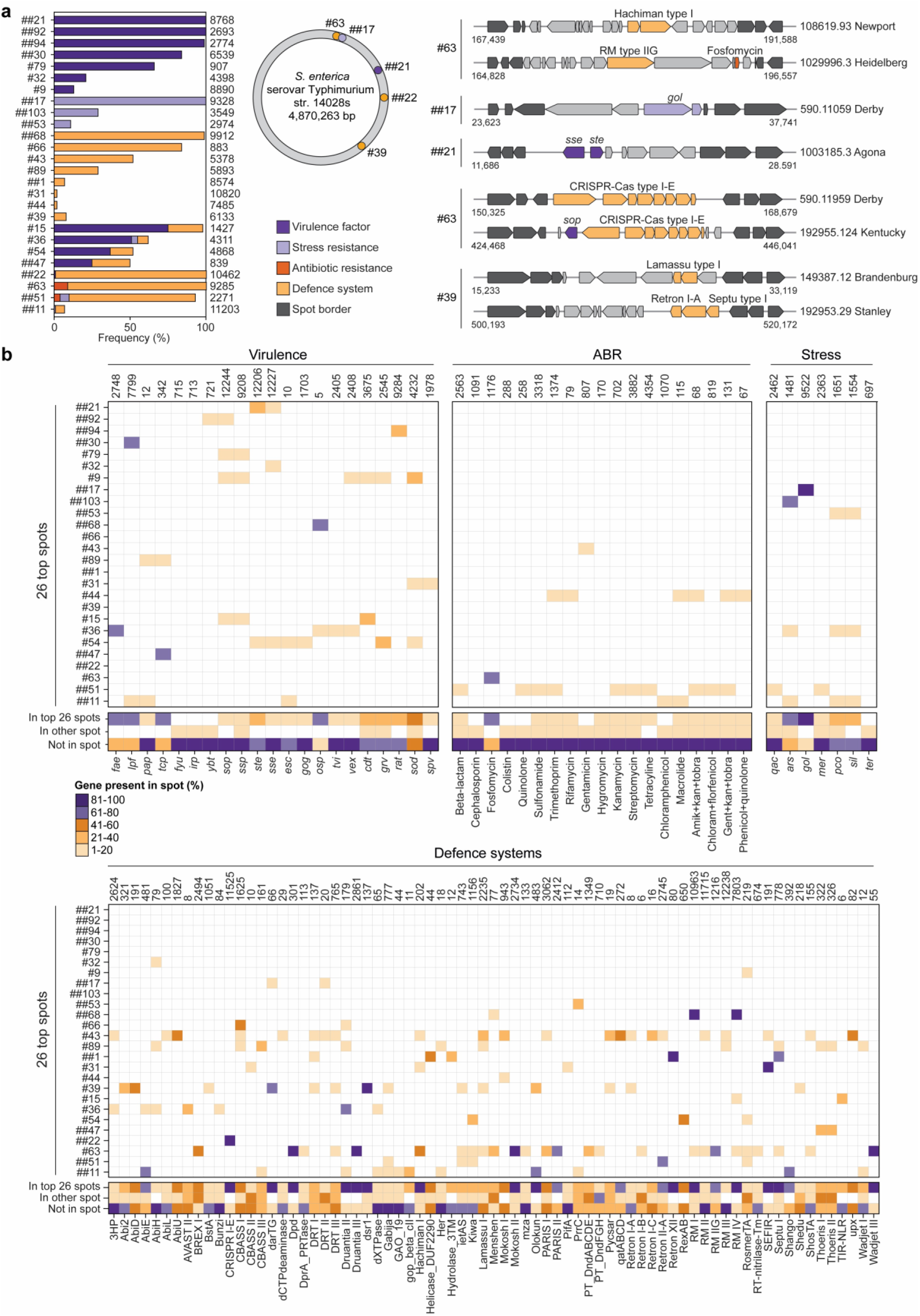
| Distribution of gene content in the spots of *Salmonella* genomes. **a** Distribution frequency of pathogenicity determinants across spots identified in at least 1% strains of *Salmonella.* Mapping of spots present on the reference strain *S. enterica* subsp. *enterica* serovar Typhimurium str. 14028s, with examples illustrating various gene arrangements. **b** Frequency at which a specific gene cluster appears outside a spot, in one of the 26 most abundant spots, or in other spots. Only gene clusters found in spots are depicted.

Another important example of gene cluster preference for distinct spots involves the type I-E CRISPR-Cas system, predominantly found in spot ##22. Spot ##22 is ubiquitously present across all species except for serovars Brandenburg, Java, Javiana, Johannesburg, Lubbock, Mbandaka, Panama, Reading, and Worthington (**Supplementary Table 12**). In these serovars, the CRISPR-Cas system is either entirely absent or present in only a limited number of strains (**Supplementary** Fig. 1). The well-conserved nature of the type I-E CRISPR-Cas system in *Salmonella*^54,55^ seems intrinsically tied to the widespread prevalence of spot ##22 (90% of all strains). Additionally, RM types I and IV have a strong preference for spot ##68, which is present in over 80% of the strains, thus accounting for the wide prevalence of RM systems in *Salmonella*.

On a broader scale, we also observe a general inclination for gene clusters with related functions to cluster within the same spots. This is well demonstrated by particular spots housing diverse anti-phage defences, functioning as hotspots for variable defence systems (e.g., ##11, #39, #43, and #63). For example, spot #63 contains an array of defence systems, with Dpd, Druantia type III, Mokosh type II, and Wadjet type III present in more than 80% of instances (**Fig. 4a,b**, **Supplementary Table 12**).

In summary, our findings reveal the preference of gene clusters to integrate into specific spots within the *Salmonella* genome, with the presence of these spots indicating the potential presence of these gene clusters. These dynamic regions play a critical role in bacterial adaptability and fitness, as evidenced by the exclusive association of the type I-E CRISPR-Cas system with serovars containing spot ##22.

### Spot flanking genes are likely determinants of gene cluster preference

We investigated whether the preference of the gene clusters for particular spots was influenced by the gene neighbourhood, particularly the highly conserved genes flanking the spots. To accomplish this, we examined the genomic locations of the 26 prevalent spots identified in our study (an interactive visualisation of the gene content of the spots can be found at the associated Github, see Data Availability).

While some flanking genes have unknown functions, their predicted roles suggest potential connections to spot functionality. For instance, spots ##1 and #63, which seem to be preferred by anti-phage defence systems, are associated with helix-turn-helix (HTH)-type transcriptional regulators (*ecpR* and *gntR*, respectively) as border genes (**Supplementary Table 13**). Gene *ecpR* in spot ##1 is negatively regulated by H-NS and positively regulated by itself and integration host factor (IHF), a protein involved in various phage-related processes such as integration and propagation^56^. The regulatory interactions involving *ecpR* and IHF suggest a potential influence of the first on phage-related processes. Similarly, spot #63 features the flanking gene *gntR*, which influences cell wall permeability and bacterial motility, both factors known to affect phage infectivity^57^. Additionally, spot #63 contains a flanking gene encoding protein L-threonine 3-dehydrogenase, and L-threonine has been observed to impact phage infection in *E. coli*^58^. Spot ##22 houses the CRISPR-Cas type I-E defence system, and is adjacent to the *cysD* gene involved in cysteine biosynthesis^59^. Notably, the regulation of the *cysD* gene involves the *cnpB* gene, which participates also in CRISPR-Cas regulation^60^. This co-localisation suggests potential coordination between these genes for regulatory purposes.

Spot ##17 is a hotspot for gold resistance and is flanked by the *oprM* gene. *oprM* encodes an outer membrane protein that functions as an antibiotic and metal pump^61^, enabling the bacterium to defend against the toxicity of antibiotics and metals. This suggests that the presence of *oprM* in the vicinity of spot ##17 contributes to gold resistance by facilitating the efflux of gold ions from the bacterial cell. Similarly, spot ##53, associated with copper and silver resistance genes, is adjacent to the *uspB* gene encoding a universal stress response protein^62^. UspB promotes cell survival and protects against stress-induced damage, potentially aiding the bacterium in coping with copper and silver stress.

Spot ##21 contains the *ste* and *see* genes involved in *Salmonella* enterotoxin production, and is flanked by the *mdoD* gene, which encodes glucan biosynthesis protein D. This protein is essential for the synthesis of osmoregulated periplasmic glucans, which contribute to the stability and integrity of the bacterial cell envelope^63^. Although no direct connection between periplasmic glucans and enterotoxin production has been reported, it is possible that glucan production and the secretion systems responsible for enterotoxin export indirectly influence each other through broader cellular processes or regulatory networks. Spot ##30 contains the *lpf* gene cluster responsible for the production of long polar fimbriae, which facilitates bacterial adherence and colonisation of host cells and tissues^64,65^. This spot is flanked by the *eptB* gene encoding Kdo(2)-lipid A phosphoethanolamine 7’’-transferase, an enzyme that modifies the lipopolysaccharide (LPS) lipid A portion, contributing to bacterial resistance against cationic antimicrobial peptides^66^. *eptB* may support the survival of *Salmonella* adhering to host cells via *lpf* by protecting the bacterial cells from the antimicrobial peptides produced by endothelial cells, particularly in environments like the gastrointestinal tract.

Finally, spot ##51, which harbours several antibiotic resistance genes, is flanked by the *yidC* and *mdtL* genes. The *yidC* gene encodes the membrane protein YidC, crucial for the insertion and folding of membrane proteins in bacteria. YidC is implicated in the proper folding and assembly of essential membrane proteins associated with antibiotic resistance mechanisms and has been proposed as a potential antibiotic target^67,68^. On the other hand, the *mdtL* gene encodes a multidrug efflux transporter protein responsible for exporting a wide range of drugs and toxic compounds out of the bacterial cell, contributing to antibiotic resistance.

In conclusion, our investigation revealed potential relationships between spot functionality and the genes in their vicinity. The identification of specific flanking genes suggests their involvement in various processes related to phage defence, metal resistance, stress response, and antibiotic resistance. These spatial arrangements provide insights into potential coordination, regulatory connections, and adaptive mechanisms within bacterial genomes.

## Discussion

The genetic landscape of *Salmonella* is a mosaic shaped by various factors, with RGP acting as significant contributors. These dynamic genomic segments house diverse gene clusters that hold the potential to dictate the genetic makeup of *Salmonella* strains. In the traditional view, gene distribution within a genome was often perceived as a stochastic process, but recent insights have challenged this view by revealing that genes linked to specific functions tend to cluster in certain regions^69,70^. The extent and implications of this phenomenon for bacterial evolution and adaptation have remained largely unexplored.

Our findings revealed a distinctive pattern of non-random integration of gene clusters into specific RGP of *Salmonella*. Exploring the prevalence of certain RGP across diverse lineages of *Salmonella* revealed their pivotal role in shaping genetic content, and thus the pathogenicity and survival strategies of each lineage. Noteworthy examples include the presence of SPI-2 in one strain of *S. bongori*, indicating it may have developed the ability to infect warm blooded hosts. This divergence challenges established notions of gene distribution and exemplifies how RGP can redefine our understanding of gene presence and absence across lineages. Furthermore, the association between the absence of type I-E CRISPR-Cas systems and the lack of spot #22 provides a rationale for the previous observation of the missing type I-E CRISPR-Cas system in specific *Salmonella* serovars^71^.

The mobility of gene clusters across genomes has raised questions about their integration without guaranteed regulation of expression upon insertion^72–75^. Potential problems include situations where the cluster relies on regulatory interactions that are not present in the new host, genes fail to express correctly, or auxiliary interactions and dependencies on the host come into play^76^. However, our study demonstrates a non-random integration pattern of RGP and their associated gene clusters, suggesting a purposeful selection of locations rather than randomness. Bacterial genomes are often organised in gene clusters regulated by shared regulators^77^, supporting the idea that RGP are placed in specific spots primarily due to the benefit of co-regulation. This suggests that genes flanking certain genomic spots might dictate the integration of particular RGP. For instance, the strategic insertion of genes responsible for long polar fimbriae production in regions flanked by antimicrobial peptide resistance genes suggests functional synergy, potentially aiding survival of *Salmonella* by protecting it from host-produced antimicrobial peptides during invasion. This underscores the likely coordination of gene expression among co-localised genes. Similarly, the positioning of stress resistance genes near specific RGP, harbouring metal resistance genes (##17, #53) might reflect a fine-tuned regulatory network to efficiently counter stressors. Notably, hotspots associated with antibiotic resistance genes (e.g. #51) are flanked by genes implicated in antibiotic resistance mechanisms, while certain spots with anti-phage defence systems (#1, #63) are flanked by HTH transcriptional regulators linked to phage-related processes. These preferences for genomic locations might be driven by selective pressures or other factors that ensure co-expression and coordinated functionality, contributing to the intricate landscape of bacterial adaptation and evolution. Examining these potential functional links can unveil novel pathogenicity traits and gene interaction networks crucial for understanding *Salmonella* pathogenicity.

Unsurprisingly, our findings underscored the prominent role of plasmids in influencing ABR patterns within *Salmonella*. Notably, a substantial proportion (78%) of ABR genes were housed within these MGEs, particularly those of the IncN, IncA/C, IncHI1 and IncI1 plasmid groups. Noteworthy variations were observed across subspecies and serovars, with ABR genes predominantly concentrated in *S. enterica* subsp. *enterica*, raising concerns over the potential amplification of ABR due to human antibiotic usage. An especially troubling discovery was the presence of colistin resistance genes within serovars associated with human outbreaks, such as Saintpaul, Cholerasuis, and Paratyphi B. Colistin, designated as a crucial antibiotic by the World Health Organization^78^, serves as a last-resort defence against life-threatening infections caused by multidrug-resistant Gram-negative bacteria^78^. The occurrence of plasmid-borne colistin resistance within these outbreak-causing serovars^79^ carries the risk of propagation to other bacteria, including those with substantial clinical relevance.

But the role of plasmids extends beyond ABR. The IncF group favours virulence factors, aligning with previous reports^80^. Plasmids from the IncI1 and IncA/C groups are key vectors for *qac* gene dissemination, associated with antimicrobial and biocide resistance, and also carry mercury and tellurite resistance determinants. Plasmids affiliated with the IncHI2 group and of *Shigella flexneri* preferentially carry stress resistance determinants (*qac, mer, sil* and *ter*), while those from the pBBSB1 and IncHI groups emerge as prominent bearers of defence systems. The differential prevalence of these traits in *Salmonella* can be attributed to the distinct plasmid types prevalent in each lineage. For instance, IncA/C is mostly found in *S*. Typhimurium and confers resistance against mercury, tetracycline and sulfonamide, while IncHI1 and IncN, found primarily in *S*. Typhi, exhibit resistance against tetracycline and beta-lactams^81^.

Our study also reveals the involvement of prophages in contributing to pathogenicity-associated gene patterns within *Salmonella*, especially in the case of anti-phage defence systems. Moreover, specific phage genera are linked to the dissemination of other factors, with *Brunovirus* and *Peduovirus* frequently carrying virulence genes, *Traversvirus* carrying stress resistance genes, and *Xuanwuvirus* harbouring some ABR genes. Overall, our findings underscore the multifaceted contributions of plasmids and prophages in shaping the pathogenicity of diverse *Salmonella* lineages.

In the broader context, these findings offer a novel perspective on deciphering the evolutionary trajectory of *Salmonella*. By uncovering the complex relationships between gene clusters, RGP, and pathogenic attributes, we gain deeper insight into mechanisms driving the emergence of diverse *Salmonella* lineages. This knowledge not only enriches our understanding of the evolution of *Salmonella* but also holds promise for predicting its future adaptations and developing targeted interventions to combat infections.

## Methods

### Data collection

A total of 16,506 *Salmonella* genomes were downloaded from the PathoSystems Resource Integration Center (PATRIC)^82^ and NCBI genome databases in May 2021. Duplicate entries from both databases were removed. The completeness of the genome assemblies was assessed using BUSCO^83^, and strains with a recommended quality score of 95 or higher were selected. ANI scores were calculated using FastANI^84^, comparing each strain against the reference genome of *Salmonella* (*S. enterica* subsp. *enterica* serovar Typhimurium str. LT2, accession nr. CP060507.1). Strains with an ANI score of 90% or higher were retained^85^. After applying these filters, the final dataset consisted of 12,244 genomes. The MASH tool^86,87^ was used to calculate the mash distance between these strains, with a threshold of 0.1. The serovar identification and country of isolation for each strain were obtained from the information available in the PATRIC and NCBI databases. The host-specificity of the species, sub-species, or strains was determined through an extensive literature review^88–91^, categorising them as host-specific (human or poultry specific), host-adapted, or broad host range. Serovars with insufficient details were categorised as having an unknown host range.

### Phylogeny building and pangenome analysis

The genomes were clustered using the K-mer-based tool PopPUNK v2.5.0^92^. The model for *Salmonella* was fitted using dbscan and the phylogeny was visualised using Microreact^93^ and iTOL^94^. The pangenome analysis was performed using PPanGGOLin v1.2.74^46^, and the pangenome graph was visualised using Gephi software (https://gephi.org) with the ForceAtlas2 algorithm. The RGP and the spots of insertion were extracted using the panRGP^47^ subcommand of PPanGGOLin. RGP without a corresponding spot were excluded from further analysis. These included RGP on a contig border (i.e., likely incomplete) and instances in which the RGP is an entire contig (e.g. a plasmid, a region flanked with repeat sequences, or a contaminant). The frequency of the spot border gene and genes belonging to RGP were calculated using custom Python scripts (see Data Availability).

### Genome annotation and detection of genes of interest

The genomes were annotated using Prokka v1.14.6^95^. The virulence factors, antibiotic resistance, and stress resistance genes were identified using Abricate v1.0.1 (https://github.com/tseemann/abricate) against the Comprehensive Antibiotic Resistance Database (CARD)^96^, NCBI AMRFinderPlus^97^ and Virulence Factor Database (VFDB)^98^. The defence systems in the genomes were identified using PADLOC v1.1.0^99^ and DefenseFinder v1.0.9^100^. The genes classified as adaptation or other categories were removed. Duplicate hits with the same gene name and location were removed using custom Python scripts (see Data Availability). Quorum-sensing genes were detected using the automatic annotation process of QSP v1.0 (https://github.com/chunxiao-dcx/QSAP) from the QS-related protein (QSP) database^101^. The virulence factors, antibiotic resistance genes, stress resistance genes, and defence systems were considered to be part of a particular RGP if the entire system was within the RGP.

### Detection of plasmid, prophage and mobilome

Platon v1.6^102^, with default settings, was used to detect and annotate plasmids in the assemblies. Plasmid PubMLST^103^ was used for plasmid typing to determine the incompatibility groups. The *Salmonella* plasmid virulence (spv) region was identified by referencing the VFDB database and mapped onto the plasmid contigs to identify pSV. A representative pSV was visualised using Geneious Prime v2022.2.2. Prophage regions were identified using Phigaro v2.2.6^104^ on default mode and PhageBoost v0.1.3^105^ with a score >0.7 and a subsequent filtering with Phager (https://phager.ku.dk). From phages identified, duplicates were removed using Dedupe (https://github.com/dedupeio/dedupe) with a minimum identity of 100% and clustered at 95% identity across the region. taxmyPHAGE (https://github.com/amillard/tax_myPHAGE) was run on these regions to identify the phage kingdom, phylum, class, genus, species and name. For virulence factors, antibiotic or stress resistance genes, or defence systems to be considered within the prophage, the entire gene cluster had to be located within the prophage region. Heat maps were generated using GraphPad Prism v9.2.0.

### Statistical analyses

Statistical analyses were performed using GraphPad Prism v9.2.0, employing simple linear regression and Pearson correlation analysis with a significance level set at a two-tailed P-value with a confidence interval of 95% for the correlation between the count of plasmid and antibiotic resistance genes.

## Data availability

All original code has been deposited at GitHub, https://simrankushwaha.github.io/Genome-Plasticity-in-Salmonella/. An interactive visualisation of the gene content of the spots is also available at GitHub. The metadata of the isolates, phylogenetic analysis and the country of isolation can be visualised on Microreact, https://microreact.org/project/pRbGPKfTYKTHJfDipWbZze-project1-genomic-plasticity-is-a-blueprint-of-diversity-in-salmonella-lineages and https://microreact.org/project/mpAsGwTnnkzzFqQoFLP8g3-project2-genomic-plasticity-is-a-blueprint-of-diversity-in-salmonella-lineages.

## Acknowledgements

We acknowledge the use of the IRIDIS High-Performance Computing Facility at the University of Southampton. This work was supported by the British Council Newton Bhabha Fund [grant number 654669088] to S.K.K. and Wessex Medical Trust [grant number AB03] to F.L.N. Infrastructure at the Center for Evolutionary Hologenomics was funded by the Danish National Research Foundation grant DNRF143.

## Author contributions

Conceptualisation, F.L.N.; Methodology, S.K.K. and F.L.N.; Software: S.K.K., A.A., Y.W., H.L.A., T.S.P.; Formal Analysis S.K.K., S.A.M., A.M., and F.L.N.; Investigation, S.K.K., A.A., H.L.A., Y.W.; Data Curation, S.K.K and F.L.N.; Writing – Original Draft, S.K.K. and F.L.N.; Writing – Review & Editing, all authors; Supervision, S.A.M., and F.L.N.; Funding Acquisition, S.K.K. and F.L.N.

## Supplementary information

**Supplementary Fig. 1 | Distribution of virulence factors, antibiotic resistance (ABR) genes, stress resistance genes, and defence system across *Salmonella***. The prevalence of the specific gene (cluster) in chromosome, prophage, or plasmid is shown as a bar graph.

**Supplementary Fig. 2 | Distribution of pathogenicity determinants on plasmid and prophage classes. a** Correlation between number of plasmids and number of ABR genes in *Salmonella* strains. **b** Frequency distribution of virulence factors, antibiotic resistance (ABR) genes, stress resistance genes, and defence systems on different prophage genera. **c** Frequency distribution of virulence factors, ABR genes, stress resistance genes, and defence systems on different plasmid incompatibility groups.

**Supplementary Table 1.** Features of the 12,244 *Salmonella* genomes analysed in this study.

**Supplementary Table 2.** Features of the plasmids found across *Salmonella* species, sub-species, and serovars.

**Supplementary Table 3.** Prevalence of plasmid incompatibility groups across *Salmonella* species, subspecies, and serovars.

**Supplementary Table 4.** Features of the prophages found across *Salmonella* species, sub-species, and serovar.

**Supplementary Table 5.** List of pathogenicity genes analysed in this study.

**Supplementary Table 6.** Distribution and location of virulence factors, antibiotic resistance genes, stress resistance genes, and defence systems in *Salmonella* species, sub-species, and serovars.

**Supplementary Table 7.** Average of plasmids, prophages, virulence factors, antibiotic resistance genes, stress resistance genes, and defence systems in *Salmonella* species, sub-species, and serovars.

**Supplementary Table 8.** Prevalence (%) of virulence factors, antibiotic resistance genes, stress resistance genes, and defence systems across plasmid incompatibility groups.

**Supplementary Table 9.** Core, persistent, shell and cloud genes identified in *Salmonella*, and their predicted function.

**Supplementary Table 10.** Regions of genomic plasticity (RGP) identified in *Salmonella* species, sub-species, and serovars.

**Supplementary Table 11.** RGP families and gene count in *Salmonella* spots.

**Supplementary Table 12.** Virulence factors, antibiotic resistance genes, stress resistance genes, and defence systems present on spots of integration in *Salmonella*.

**Supplementary Table 13.** Flanking genes defining the integration spot and their predicted function.

