## Supplementary Fig. for "Genomic plasticity is a blueprint of diversity in *Salmonella* lineages"

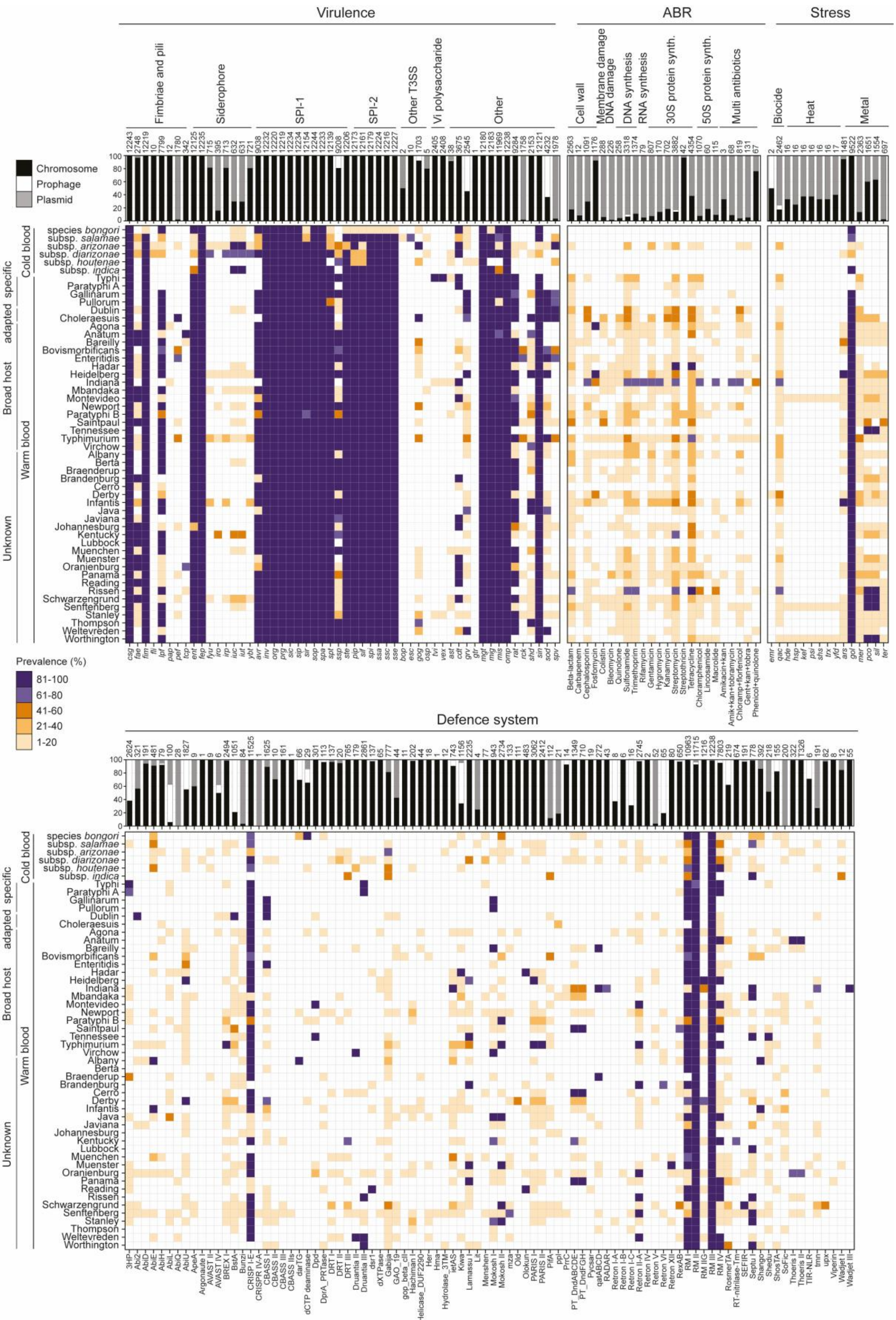


**Supplementary Fig. 1 |Distribution of virulence factors, antibiotic resistance (ABR) genes, stress resistance genes, and defence system across *Salmonella*.** The prevalence of the specific gene (cluster) in chromosome, prophage, or plasmid is shown as a bar graph.

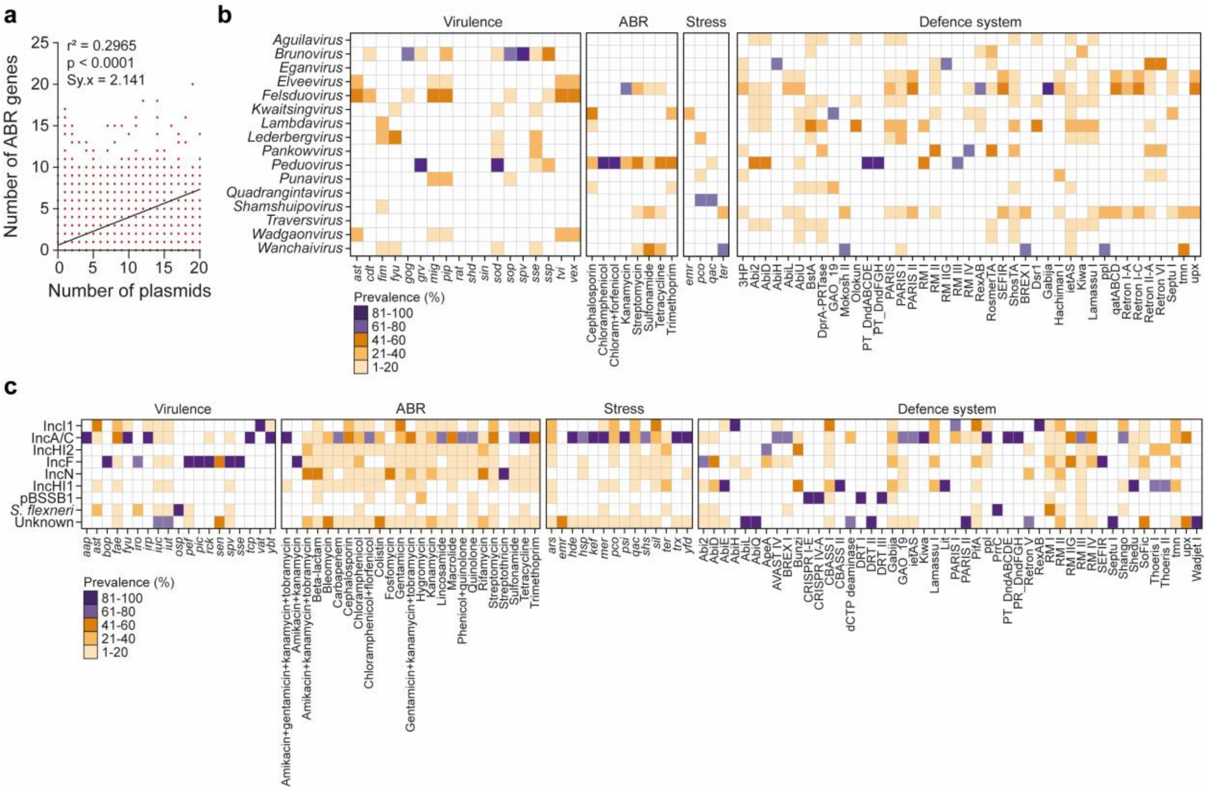


**Supplementary Fig. 2 | Distribution of pathogenicity determinants on plasmid and prophage classes. a** Correlation between number of plasmids and number of ABR genes in *Salmonella* strains. **b** Frequency distribution of virulence factors, antibiotic resistance (ABR) genes, stress resistance genes, and defence systems on different plasmid genera. **c** Frequency distribution of virulence factors, ABR genes, stress resistance genes, and defence systems on different plasmid incompatibility groups.
